# Unlocking Herbal Potentials: Novel Shikimate Kinase Inhibitors in the Fight Against Antibiotic Resistance

**DOI:** 10.1101/2024.03.02.583126

**Authors:** Siddharth Nirgudkar, Yurong Chai

**Affiliations:** Acton-Boxborough Regional High School, 36 Charter Road, Acton, 01720, MA, USA; Department of Biology, Northeastern University, 360 Huntington Avenue, Boston, 02115, MA, USA

## Abstract

Against a backdrop of stagnant antibiotic innovation, the escalating prevalence of antibiotic-resistant pathogens forecasts a challenging future [6]. Traditionally, antibiotics, predominantly derived from fungal sources, employ a limited set of mechanisms to inhibit bacterial growth [6, 16]. Shikimate Kinase has emerged as a promising antibacterial target due to its exclusivity to bacteria and the lethality of its inhibition [4, 13, 14, 15, 28, 29, 30]. Although synthetic inhibitors have been developed, the exploration of plant-derived alternatives remains untapped. Naturally derived plant-based compounds provide a more viable option because of the high cost of creating synthetic compounds. This study examines the Goldenrod plant, reputed in Native American Ethno-medicine for its antimicrobial properties [3, 12, 17]. Employing Liquid Chromatography - Mass Spectrometry (LC-MS) and Quantitative Structure Activity Relationship (QSAR) models, the study evaluates the plant’s compounds for their potential as antibacterial agents. Antibacterial activity against *Bacillus Subtilis* was assessed using the Kirby-Bauer Disk Diffusion assay, and genetic sequencing was performed on mutants that overcame the initial inhibition zone. By comparing the parent and mutant strains, the mode of inhibition by the plant antibiotic was determined by backtracking. The study identified Shikimate Kinase as the inhibitory target of the plant-derived compounds. Molecular docking revealed a binding affinity of -8.9 kcal/mol for the most effective compound, which is statistically significant compared to Shikimate Acid, the enzyme’s natural substrate. Through Pymol visualization, competitive inhibition was confirmed, with the compound’s binding pocket exhibiting a druggability score of 0.84, approaching the threshold of clinical drugs. This research suggests new antibiotic classes targeting the Shikimate Kinase pathway, offering an alternative approach to tackling ESKAPE pathogens and enhancing health outcomes.

## 1 Introduction

The World Health Organization has declared Antibiotic Winter a public crisis of unimagined levels of magnitude [6]. Despite urgently needing new antibiotics, there has been a significant antibiotic development drought since the late 80s [16]. Furthermore, any antibiotics produced since then have only had two substantial classes or ways to inhibit the growth of bacteria [16]. From this, many opportunistic pathogens have developed resistance to current antibiotics. The group of the six most nimble resistors has been labeled as the ESKAPE pathogens, a namesake for their tendency to become resistant to current antibiotics. There is a significant demand for innovative drugs and treatment modalities, especially those that operate via novel mechanisms. Additionally, a deeper insight into antibiotic interactions within a biological system is paramount.

One strategy for developing novel antibiotics is to target critical metabolic pathways unique to bacteria, allowing them to be lethal only to bacteria and not the humans they are trying to protect. The Shikimate pathway, a critical metabolic route producing chorismate, is crucial for synthesizing aromatic amino acids in bacteria, including *Bacillus subtilis* and several ESKAPE pathogens. As mammals lack this pathway, it presents an ideal target for selective antibiotic action. Specifically, the enzyme Shikimate Kinase, which facilitates a pivotal step in the pathway, stands out as a promising target for antibiotic development [24, 25]. Inhibitors of Shikimate Kinase have the potential to thwart bacterial growth by blocking the production of essential compounds derived from chorismate, thereby offering a strategic point of intervention in combating antibiotic resistance. In addition to the inhibition of chorismate production, Shikimate Kinase inhibition causes Shikimate Acid buildup, which is fatal to many bacteria (including those mentioned previously) [24, 25]. The importance of Shikimate Kinase inhibition has been studied by many through the design and synthesis of artificial drugs through crystallization projects [4, 13, 14, 15, 28, 29, 30].

For example, [28] investigates inhibitors for Mycobacterium tuberculosis Shikimate Kinase (Mt-SK). Through structural and computational studies, it explores the enzyme’s catalytic turnover, focusing on the role of conserved arginine residues in product expulsion. Substrate analogs were designed and tested, and a crystal structure of Mt-SK with potent inhibitors was solved, revealing critical insights for inhibitor design. These findings emphasize the potential of SK inhibitors as therapeutics, offering a novel approach against tuberculosis and possibly other bacterial diseases.

However, very few have looked at the natural production of these antibiotics, increasing the overall package for these antibiotics. Artificial synthesis and production of new-class antibiotics is expensive; on the other hand, having production done naturally yields much more cost-friendly for investors. In this study, *B. Subtilis* was used as a model organism due to its structural similarity in many metabolic pathways and effects to *Staphylococcus aureus*. Additionally, using plants as the source of these potential antibiotics seems to be a better option now, as the probability of finding novel antibiotics is much greater than in the much more thoroughly studied Domain of Fungi. Specifically, the Goldenrod (*Solidago*) was chosen in this study. The goldenrod plant has had numerous cites in Native American scriptures as having anti-inflammatory, antimicrobial, and even anti-cancerous properties. Despite its rich history of being used as herbal medicine, there has been very little modern research involving the goldenrod plant and its medical use, making further research on it very fruitful. The claims of the antimicrobial properties of the goldenrod plant were validated by past lab work and will now be built upon. Through the use of *in vitro* techniques as well as *in Silico* techniques, new Shikimate Inhibitors were found that were being produced naturally from the Goldenrod.

## 2 Methods

### 2.1 Preliminary Investigation

#### 2.1.1 Preparation of Plant Extracts

The goldenrod plant that grew near the Lab was chosen, and the samples were collected in late November 2022. The plant samples were stored in an incubator for seven days at 37^*°*^*C* to remove moisture, which might have contained other trace elements that could corrupt the samples. Once dried, the plant was rinsed in Distilled Water and put in the incubator to dry for 24 hours. The leaves, roots, and stems were separated and crushed individually with a coffee grinder until they were fine powder. One gram of each extract was mixed with 3ml of methanol and stored in 5ml Eppendorf tubes. Each tube was vortexed for 1 minute, steeped for 15 minutes, vortexed for 15 seconds, and allowed to steep for 1 minute. Afterward, they were centrifuged at 13,400 rpm for 1 minute. The supernatant was harvested from each and used as the final plant extract to be utilized later in the procedures.

#### 2.1.2 Cell Lines and Growth Media Preparation

The bacteria used were B.subtilis and E.coli, both standard lab grade (BSL-0). Trypsin Soy Agar (TSA) plates were used to grow E.coli and were made by using 12g of TSA powder with 100 ml of distilled water. Luria Broth (LB) plates were used to grow B.subtilis.

#### 2.1.3 Bauer-Disk Diffusion Assay

Each TSA plate was divided into five equal sections; 3 were used for the experimental variables, and two were saved for the controls. The negative control was standard 70 percent methanol, while the positive control was penicillin for B. subtilis and tetracycline for E.coli. To test the inhibitory properties, small disks were soaked in each supernatant and put on the plate; great care was taken to ensure there was no excess solution on the disks to reduce the risk of inaccurate inhibition readings. Twelve total trials were done in the preliminary investigation, six trials for each bacterium.

### 2.2 Genomic Sequencing

#### 2.2.1 Preparation of Bacterial Cultures

Due to the overall better performance of the plant to B.subtilis, it was used further to delineate the antibiotic mechanism. After plating, a preliminary zone of inhibition formed within 24 hours (Figure 4); however, after *≈* 72 hours, bacteria started to increase into the zone of inhibition, which will be labeled as MB (mutant bacteria). To understand the workings of the possible antibiotic compounds found in the goldenrod plant, each MB and the original parent bacteria (PB) was sequenced to understand the change in phenotype, leading us to possible inhibitory mechanisms. After bacterial colonies that had grown inside the preliminary inhibition zone had reached a healthy size, they were transplanted to separate cell plates and allowed to grow. Finally, these four plates (3 MB for the roots, stems, and leaves) and 1 PB were sent to the Chai Lab at Northeastern to be analyzed.

**Figure 1.**
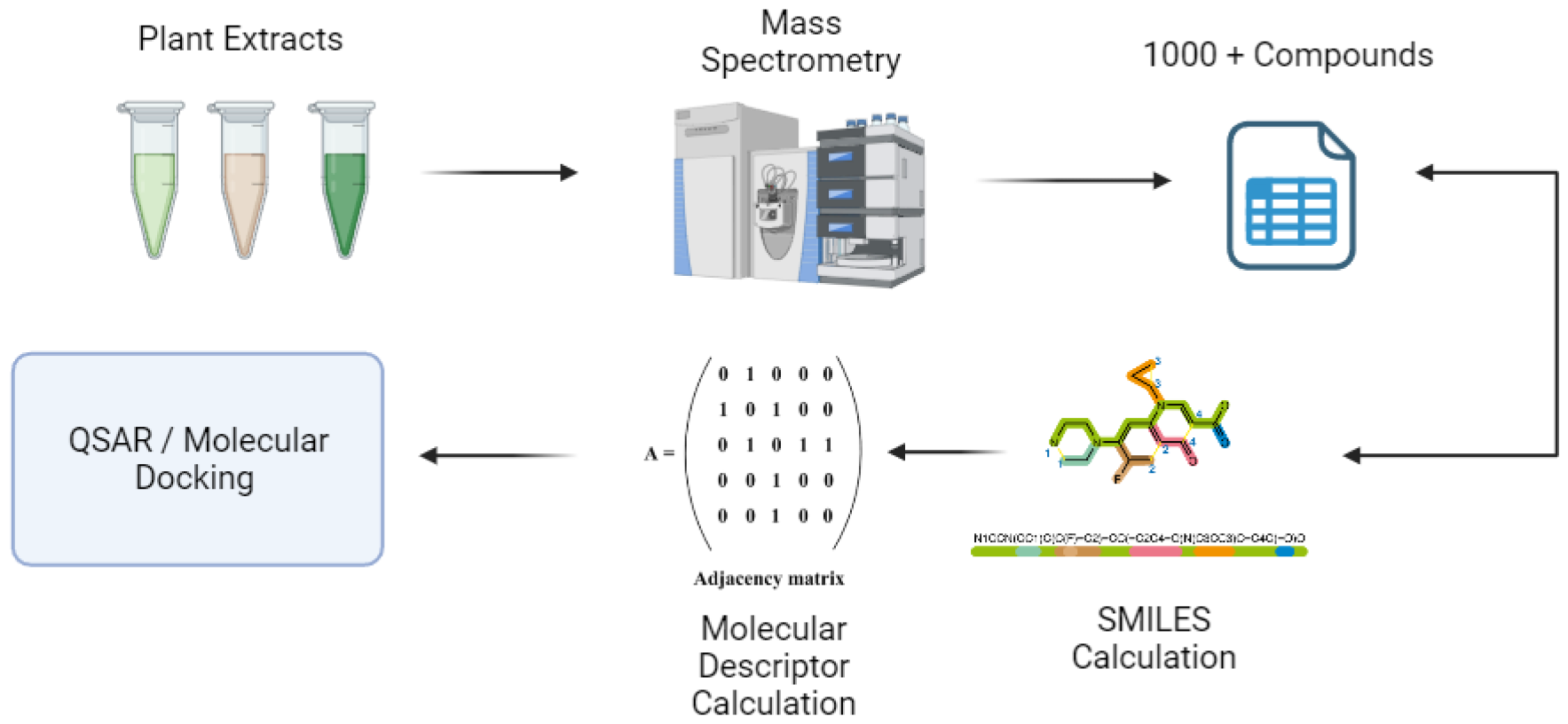
Flow chart showing the main steps in the genomic sequencing conducted in the study.

**Figure 2.**
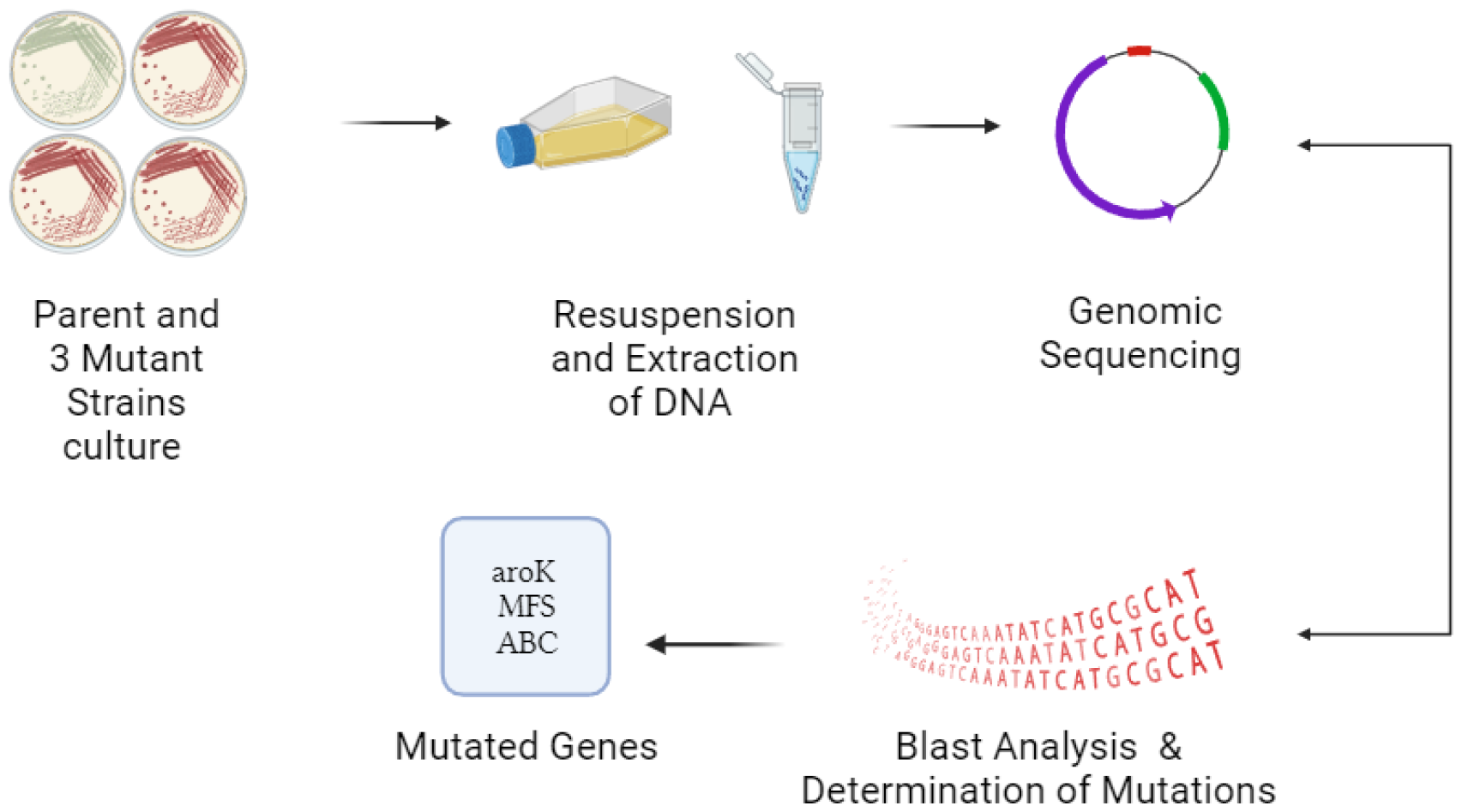
Flow chart showing the main steps involved with Compound Identification.

#### 2.2.2 Resuspension and Extraction

From the Cell Plates, a standard resuspension and extraction protocol was followed to isolate the DNA to get sequenced later.

1. Centrifuge bacterial culture in 2 mL at 8000 rpm (6800 x g) in a microcentrifuge tube for 2 minutes at room temperature. Remove all remaining medium.
2. Resuspend the cell pellet in 450 µL of autoclaved diH_2_O.
3. Add 50 µL of EDTA (0.5 M) and 60 µL of lysozyme (stock at 20 mg/mL). Invert the tubes 4-6 times. Incubate tubes at 37°C for 1 hour.
4. Add 650 µL of solution I (nuclei lysis solution) and invert the tubes 4-6 times until the solution is clear.
5. Add 250 µL of solution II (protein precipitation solution) and vortex tubes for 20 seconds.
6. Leave tubes on ice for 10 minutes, then centrifuge them at 14000xg for 5 minutes.
7. Transfer approximately 900 µL of the supernatant to a new tube and add 600 µL of isopropanol. Invert tubes 4-6 times until a white precipitate is seen.
8. Leave tubes at room temperature for 3 minutes, then centrifuge them for 5 minutes at 14000xg. Discard supernatant.
9. Wash the pellet by adding 600 µL of 70% ethanol and centrifuge tubes for 5 minutes at 14000xg.
10. Let pellet dry.
11. Resuspend pellet in 100 µL of autoclaved diH_2_O. Store DNA at -20°C.

#### 2.2.3 Genomic Sequencing

The samples were then sent to Plasmidasaurus, who performed genome sequencing. Afterward, all three mutant strains were blasted against the parent strain to determine differences in the genome. The mutations common in all three comparisons were chosen as those related to the antibiotic properties of the goldenrod plant and used further in the study. Genomic sequencing methods were derived from [10].

### 2.3 Compound Identification

#### 2.3.1 Mass Spectrometry

Following the preliminary investigation, the three samples from the plant (stems, roots, leaves) were sent to the Sjoelund Lab at Northeastern to undergo mass spectrometry. Specifically, an Orbitrap Explonis 240 Mass Spectrometer was used to detect novel compounds. Over 1000 compounds were discovered initially; however, the compounds with a confidence level of less than 0.7 were discarded, leaving 100 compounds.

#### 2.3.2 Canonical Smiles Discovery for each Compound

A Canonical smile is a unique SMILES representation of a molecule, no matter the orientation of the molecule [7]. These are especially important in drug discovery, as the calculation of molecular descriptors changes based on the smiles; the molecular descriptors are then used further to predict the properties of molecules. The QSAR analysis outputted each compound’s molecular formula and a structural picture. Many compounds’ SMILES were found using the software OSPIN [22], while others had to be hand calculated by drawing the molecules in Pub Chem’s smile detection software.

### 2.4 Quantitative structure-activity relationship (QSAR) development

#### 2.4.1 Choosing Molecular Descriptors

In developing Quantitative Structure-Activity Relationship (QSAR) models, selecting molecular descriptors is of paramount importance [5]. These models’ effectiveness hinges on using unique and carefully chosen descriptors [5]. An optimal set of descriptors should capture the essential molecular characteristics relevant to the study while avoiding redundancy. It is crucial to balance the Number of descriptors employed; an excessive number can lead to overfitting and deterioration of the model’s predictive power. Therefore, an informed choice of distinct and relevant molecular descriptors is fundamental to the robustness and accuracy of QSAR models. Because of this importance, this study followed the advice of [8] that complied with the most used/essential descriptors used in drug discovery, as that was the main focus of this research. The molecular descriptors were chosen from RDK Mordred 1826 [23], a molecular descriptor library containing 1826 molecular descriptors; 12 were chosen. The molecular descriptors’ accuracy was validated in the experiment with the accurate confusion matrixes for all QSAR predictions.

#### 2.4.2 Calculation of Molecular Descriptors

Once the molecular descriptors had been chosen, a program was made to calculate the molecular descriptors from the canonical smiles. The program was a slight modification of [CITAR].

#### 2.4.3 QSAR Development

Multiple QSAR models were run [Figure 3] through the open source model QSAR-CO [2], each needing to be trained with their own training set.

**Figure 3.**
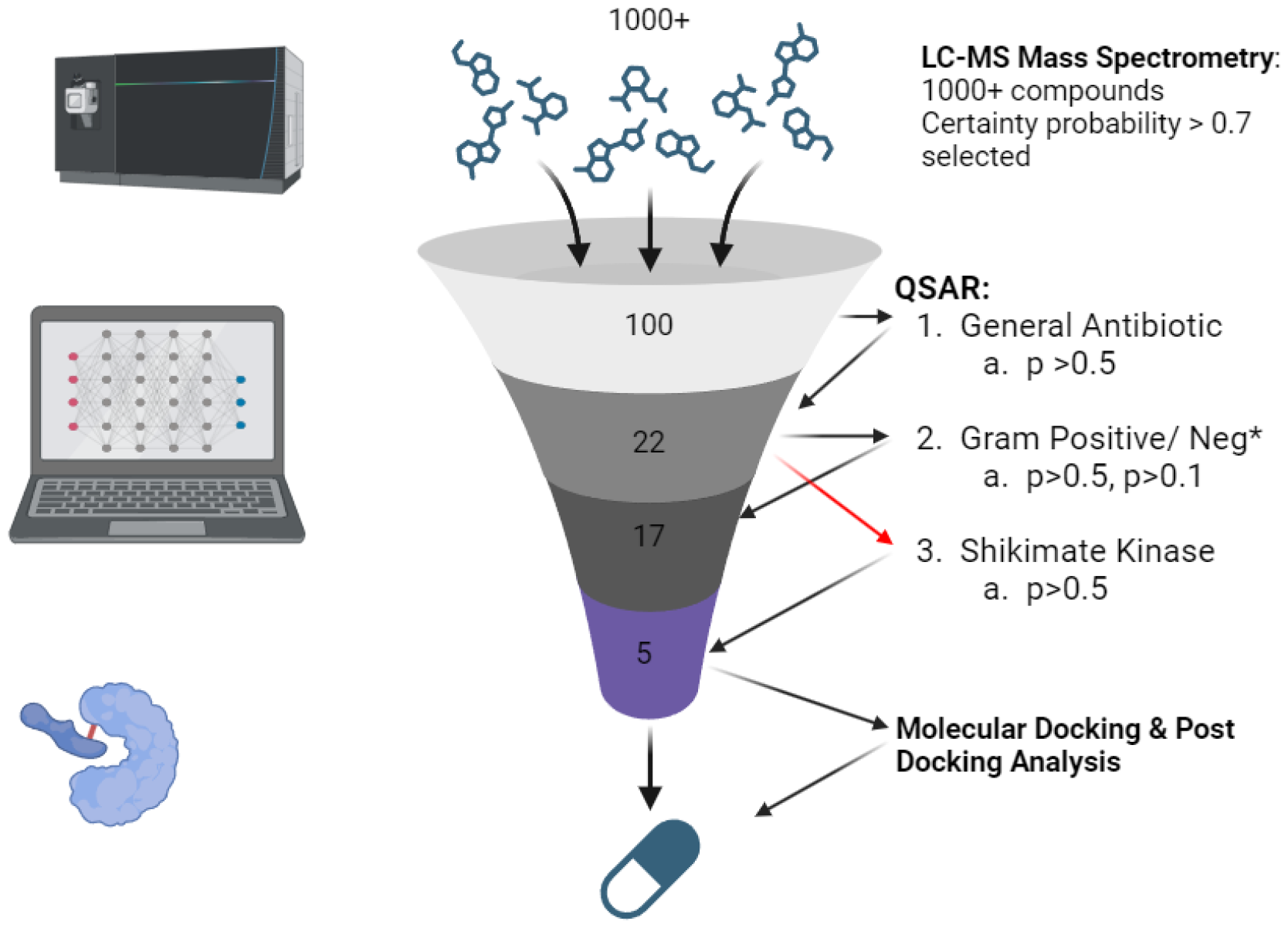
Brief Overview of the prediction process done through QSAR and Molecular Docking.

**Figure 4.**
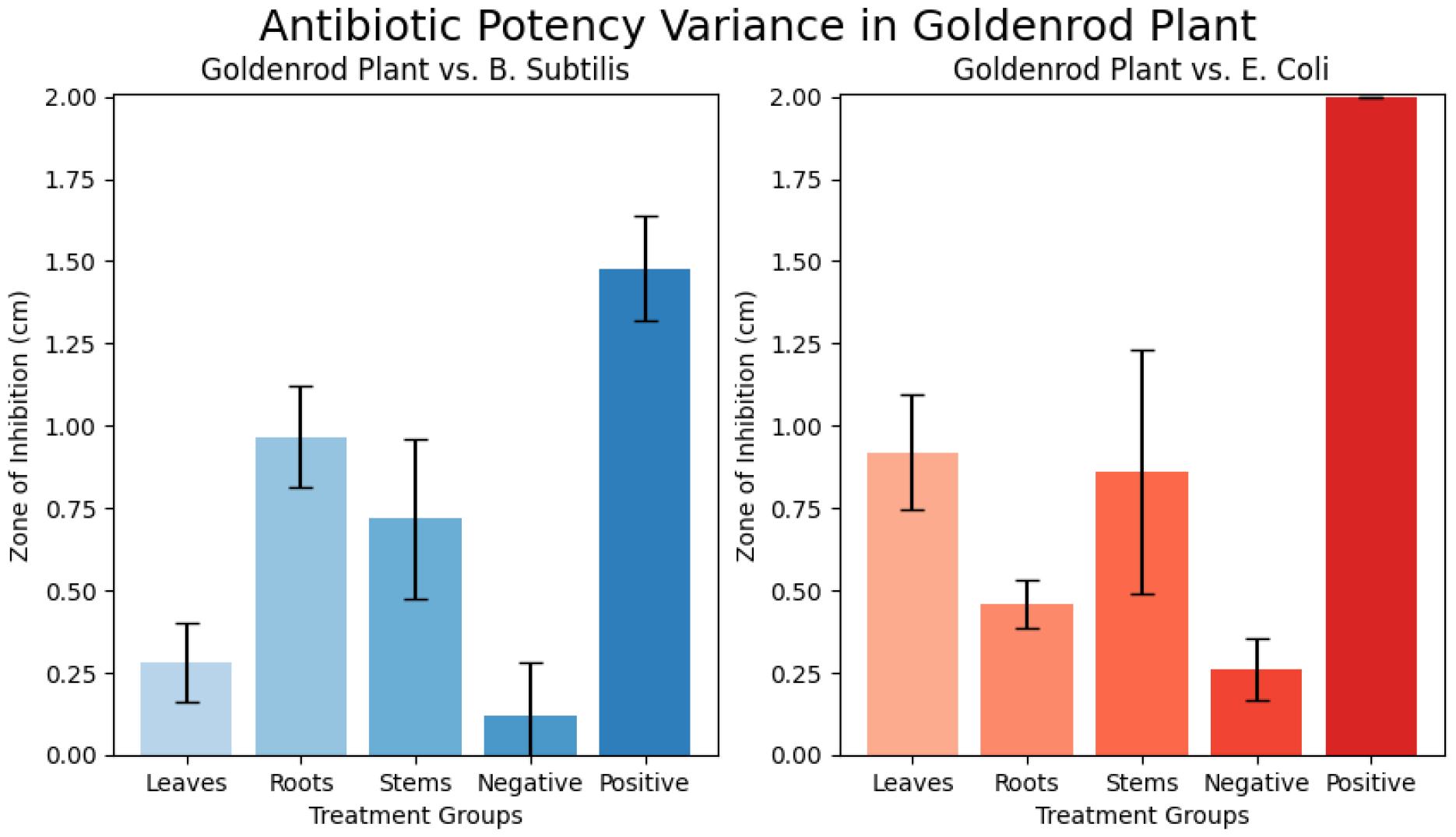
Comparative graph showing the antibiotic potency variance in the goldenrod plant against different bacteria. The black bar represents an error of 2 standard errors of the mean. The zone of inhibition was measured from the disk’s center to the nearest bacterial colony grown (following the Bauer-disk diffusion assay).

1. General Antibiotic Prediction: 102 known antibiotics were curated from the CAS database, and 73 known non-antibiotic compounds were found.
2. Gram-Positive Antibiotic Prediction: The 32 known gram-positive antibiotics were chosen (CAS), as well as 60 known non-antibiotic compounds.
3. The 70 known Gram-Negative Antibiotic compounds were chosen, and 71 known non-gram-negative antibiotic compounds were chosen
4. 11 known Shikimate Inhibitors and 30 known non-shikimate inhibitors were chosen.

Their canonical SMILES were calculated for all the chosen compounds, and then their molecular descriptors were calculated.

#### 2.4.4 QSAR Use

To finally run the QSAR, the QSAR-Co software was used [2]. The model used a normal development approach and random division to divide the training set into a true training set and a training validation (test set). The random division method was set to 70 (training) and 30 (testing) following the QSAR-Co manual. Genetic Algorithm-Linear Discriminant Analysis (GA-LDA) was chosen for QSAR model development for its efficient feature selection and capacity to manage complex datasets [32]. The genetic algorithm identifies a potent subset of descriptors, optimizing predictive accuracy, while Linear Discriminant Analysis models the descriptors’ relationship to biological activity. This strategy strikes an optimal balance between model interpretability and computational efficiency [32].

### 2.5 Molecular Docking Simulations

Once Shikimate Kinase had been identified as the target of the antibiotic compound from the goldenrod plant, molecular docking simulations were done with six ligands (compounds that the Shikimate Kinase Inhibitor QSAR predicted as positive) and the receptor (Shikiamte Kinase).

#### 2.5.1 Preparation of Molecular Files

Both the receptor and the ligands were downloaded from Pub Chem in a 3d Protein Data Bank (PDB) format for the receptor and a 3d sdf format for the ligand. The ligands were fed into Pymol and converted into pdb files [31]. The receptor was loaded into Autodock Tools, and several structural changes were made. Water molecules were excised from the crystallographic protein structure to reduce computational complexity and to avoid interference with the docking program’s intrinsic solvation model [9]. Polar hydrogens were appended to furnish a detailed representation of the protein’s hydrogen bonding landscape, ensuring the accuracy of donor-acceptor interactions in the docking predictions. Additionally, Kollmann charges were allocated to the protein’s atoms to precisely simulate the electrostatic potential, which is pivotal for the assessment of ligand-protein electrostatic congruence and the estimation of binding affinities [18].

#### 2.5.2 Use of Autodock Vina

AutoDock Vina was employed over AutoDock 4 due to its enhanced performance in terms of speed and accuracy, which are facilitated by improved scoring functions and multithreading capabilities [9, 33]. To ensure comprehensive sampling of the conformational space, an exhaustiveness parameter was set to 8, balancing computational time and the thoroughness of the search. The defined search space was a cubic box with dimensions of 40 °A on each side, centered on the active site, allowing for ample exploration of potential binding modes within a focused region of the receptor protein.

### 2.6 Post Molecular Docking Analysis

After the completion of docking simulations, a pair of computational tools were utilized to analyze the ligand-receptor interactions and to evaluate the potential druggability of the receptor’s binding pockets. Interaction profiling was conducted using the Protein-Ligand Interaction Profiler, which elucidated the nature and strength of the contacts established within the docked complexes [35]. Druggability assessment was performed using the ProteinPlus suite from Universitäat Hamburg, which assigned scores to each binding pocket on a scale from 0 to 1 [19, 34]. Binding pockets with scores of 0.7 or above were categorized as highly promising drug targets. The structural data for these analyses were sourced from the pdb formatted file of the protein-ligand complex.

## 3 Results

### 3.1 Preliminary Investigation

It has been shown in Native American literature, as well as modern findings, that the Goldenrod plant shows antimicrobial properties [3, 12, 17]. These findings were validated through a Bauer-disk diffusion assay comparing the Goldenrod plant against gram-positive and harmful bacteria, B. subtilis and E. Coli, respectively. The plant supernatant was tested against methanol (negative control), penicillin (positive control for B. subtilis), and tetracycline (positive control for gram-negative). While there is a variance between the three parts of the plant, they are all statistically more potent than the negative control; this phenomenon will be the focus of the research.

### 3.2 QSAR Experimentation

Following the mass spectrometry readings, over 1000 compounds were found, and 100 were finalized based on accuracy metrics, *p ≥* 0.7. The first trial was against known antibiotics (gram-positive, gramnegative, and broad-spectrum antibiotics). The QSAR model used a normal approach. The training set consisted of 100 known antibiotics and 74 known non-antibiotics curated from the CAS database. The QSAR model used random division, where 70 percent of the training set was used to develop the model, and 30 percent of the training set was used to validate the accuracy of the model [Figure 5]. Only the compounds that were successful predictions would move on to subsequent computational testing to avoid unnecessary computational overhead: passing the preliminary QSAR-qualified compounds as potential drugs and invalidating others.

**Figure 5.**
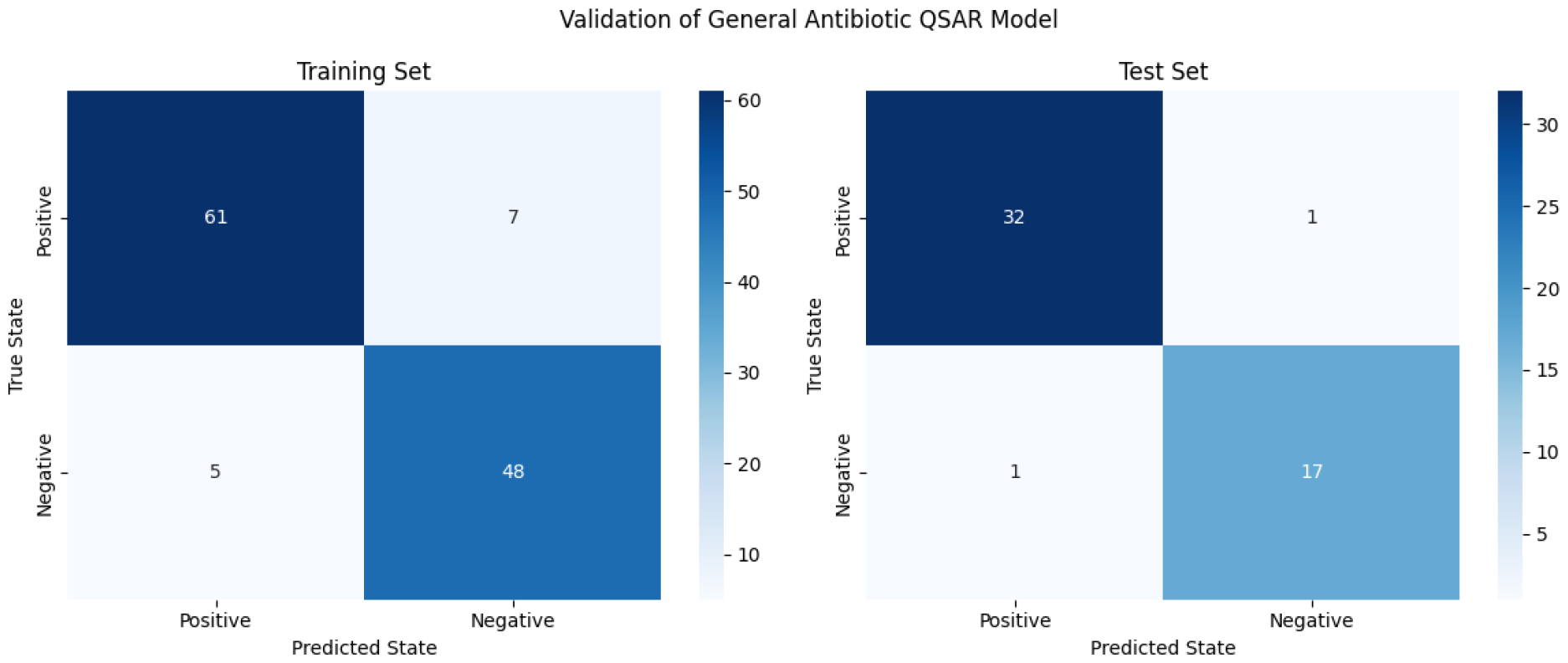
Confusion matrix for the General Antibiotic QSAR model.

**Figure 6.**
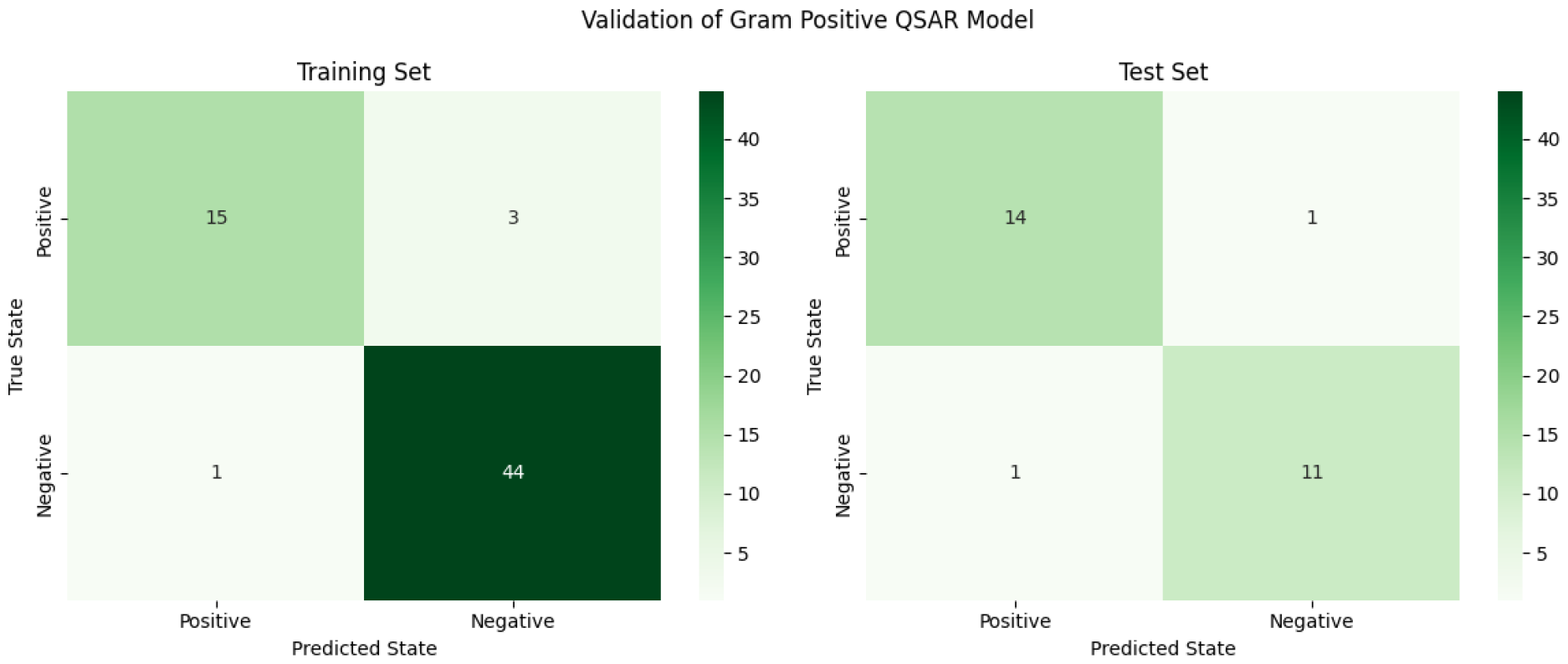
Confusion matrix for the Gram-Positive QSAR model.

**Figure 7.**
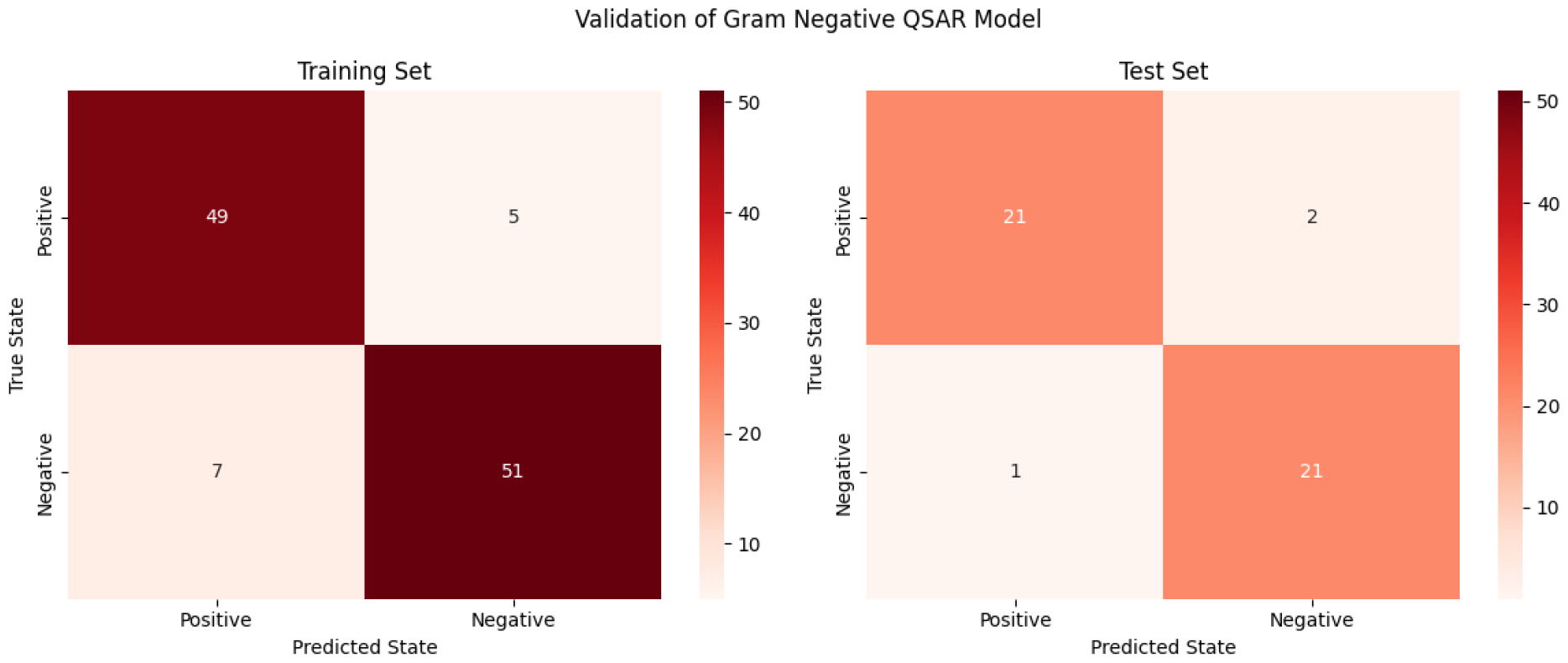
Confusion matrix of the gram-negative antibiotic QSAR model.

**Figure 8.**
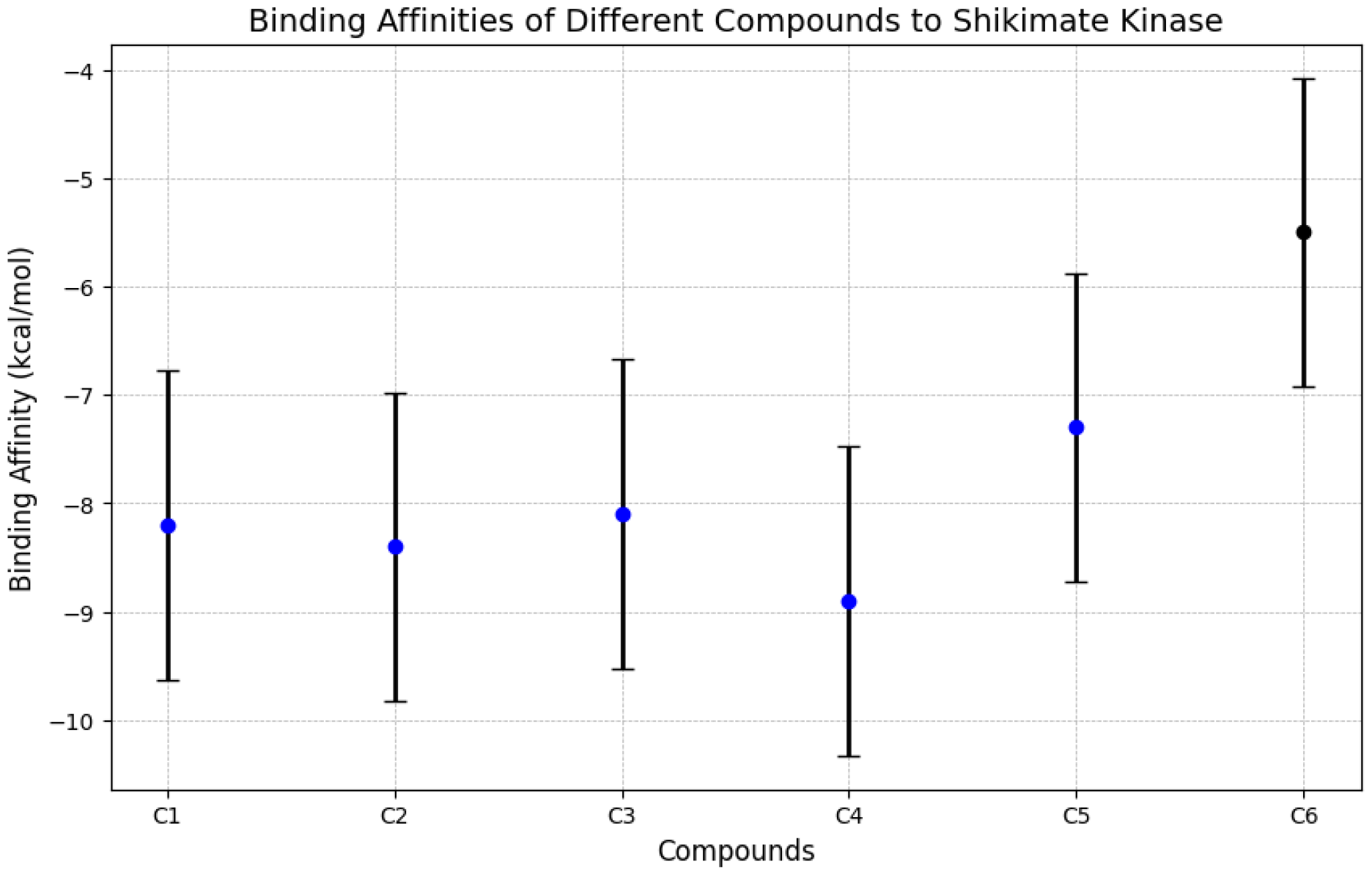
graph showing the variance in Binding Affinity kcal/mol. The compounds (C1-5) are the final compounds predicted from the QSAR model, while C6 is the natural substrate of Shikimate Kinase, shikimic acid. The black bars represent the standard error of Autodock Vina (the software used for molecular docking), which is *±* 1.42 kcal/mol. It should be noted that only compounds C2 and C4 were statistically different from Shikimic Acid, with binding affinities of -8.3 and -8.9 kcal/mol.

**Figure 9.**
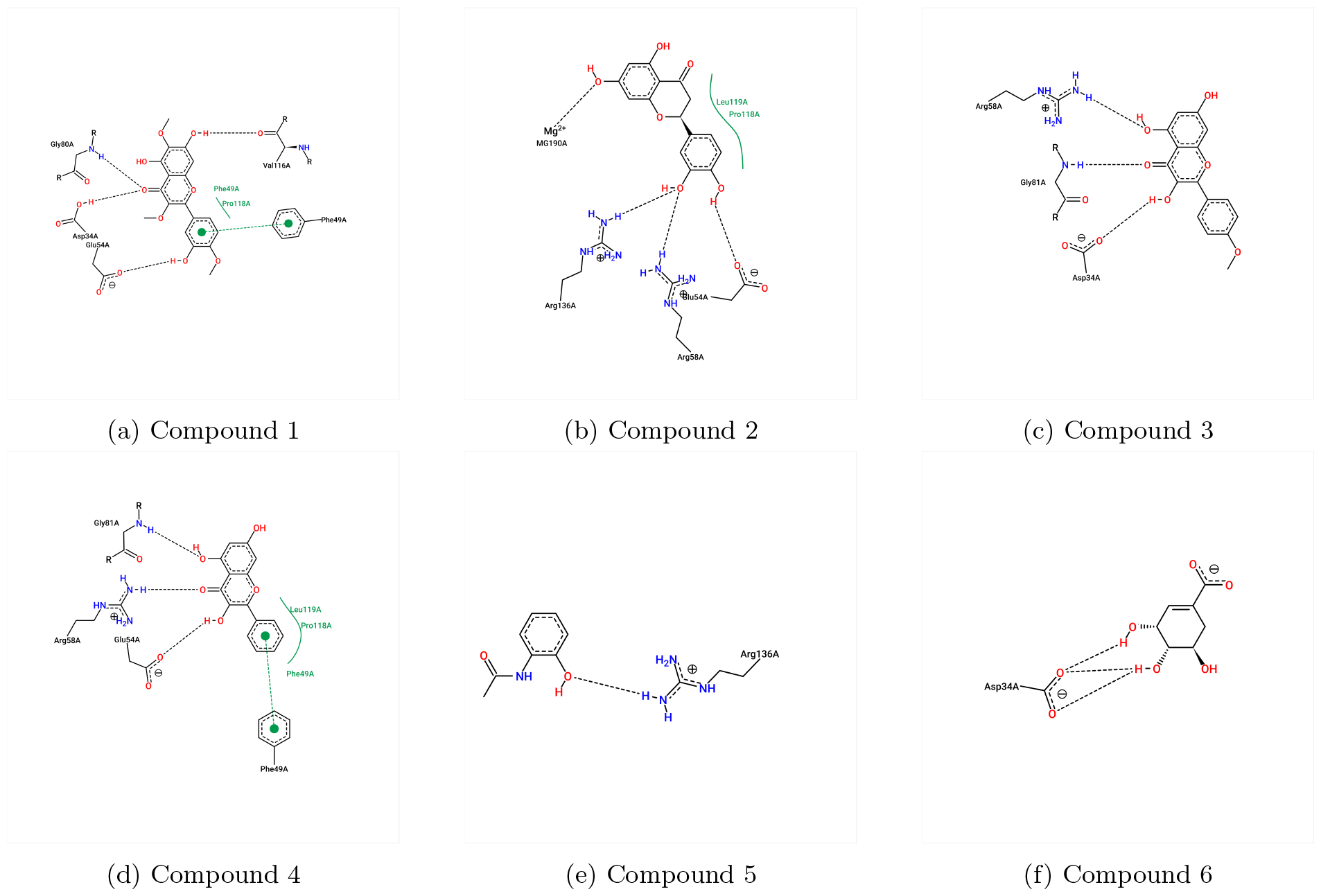
A collection of interaction diagrams between 5 possible ligands, one control (Compound 6 (Shikimate Acid)), and Shikimate Kinase.

**Figure 10.**
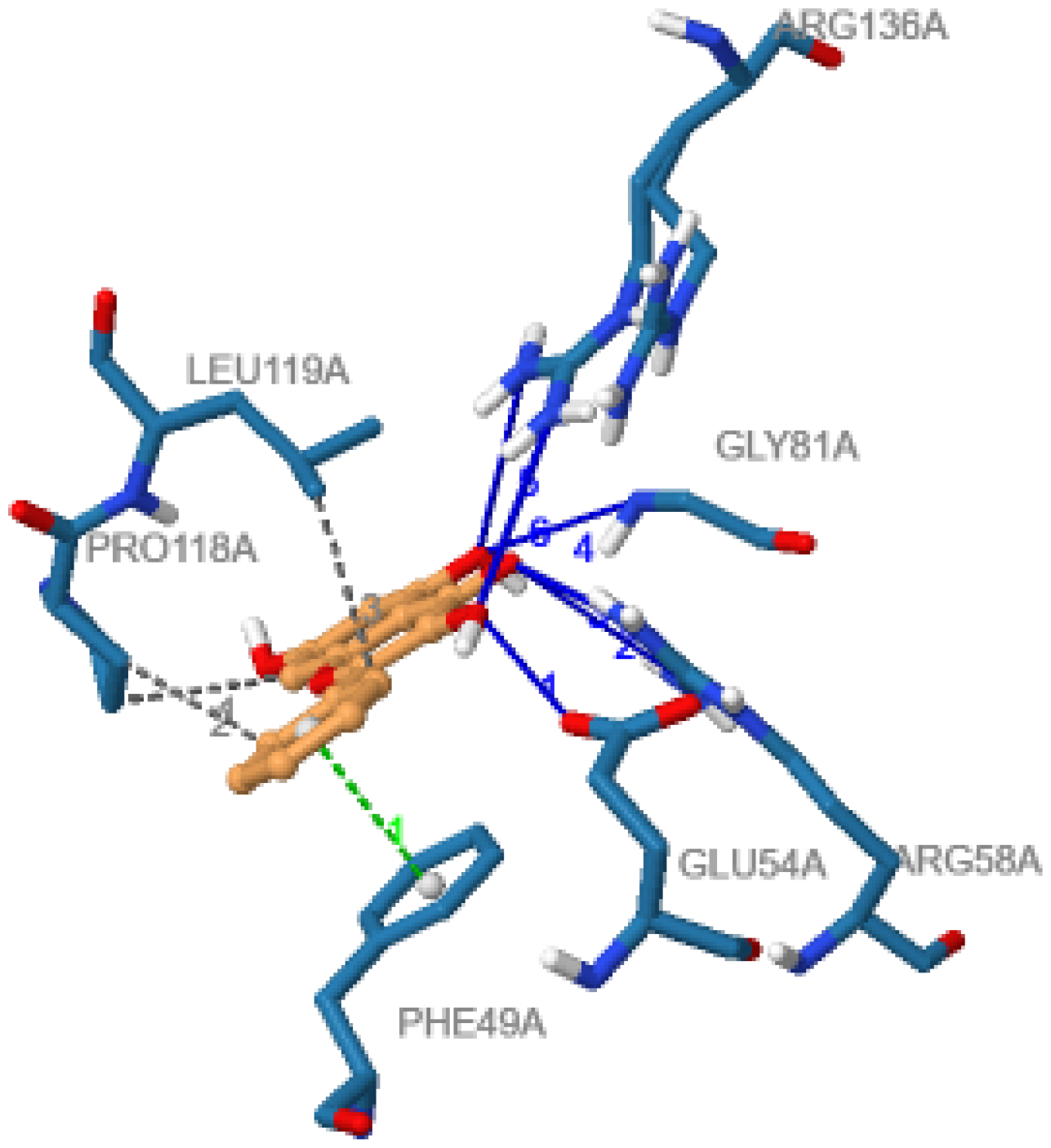
3^*d*^ Diagram showing interactions between the ligand (orange) and protein residues (blue). Grey bonds represent hydrophobic interactions, blue bonds represent hydrogen bonding, and green represents *π* stacking.

**Figure 11.**
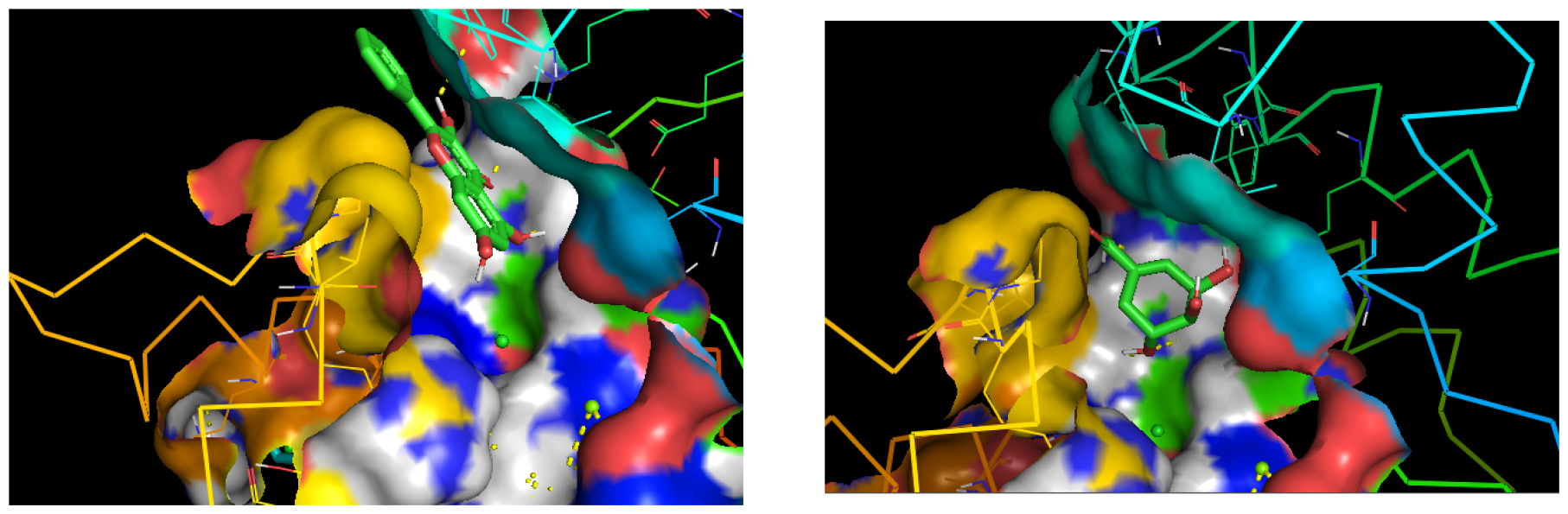
Pymol Visualization of the ligand-protein complex. C4 (Left) interacts with the same active site as Shikimate Acid (Right), showing that the type of inhibition performed by possible antibiotic C4 is most likely competitive inhibition. However, as shown in Fig. 10, interactions between ligand and amino side chains are different between C4 and C6, alluding to possible denaturation of Shikimate Kinase as well.

**Figure 12.**
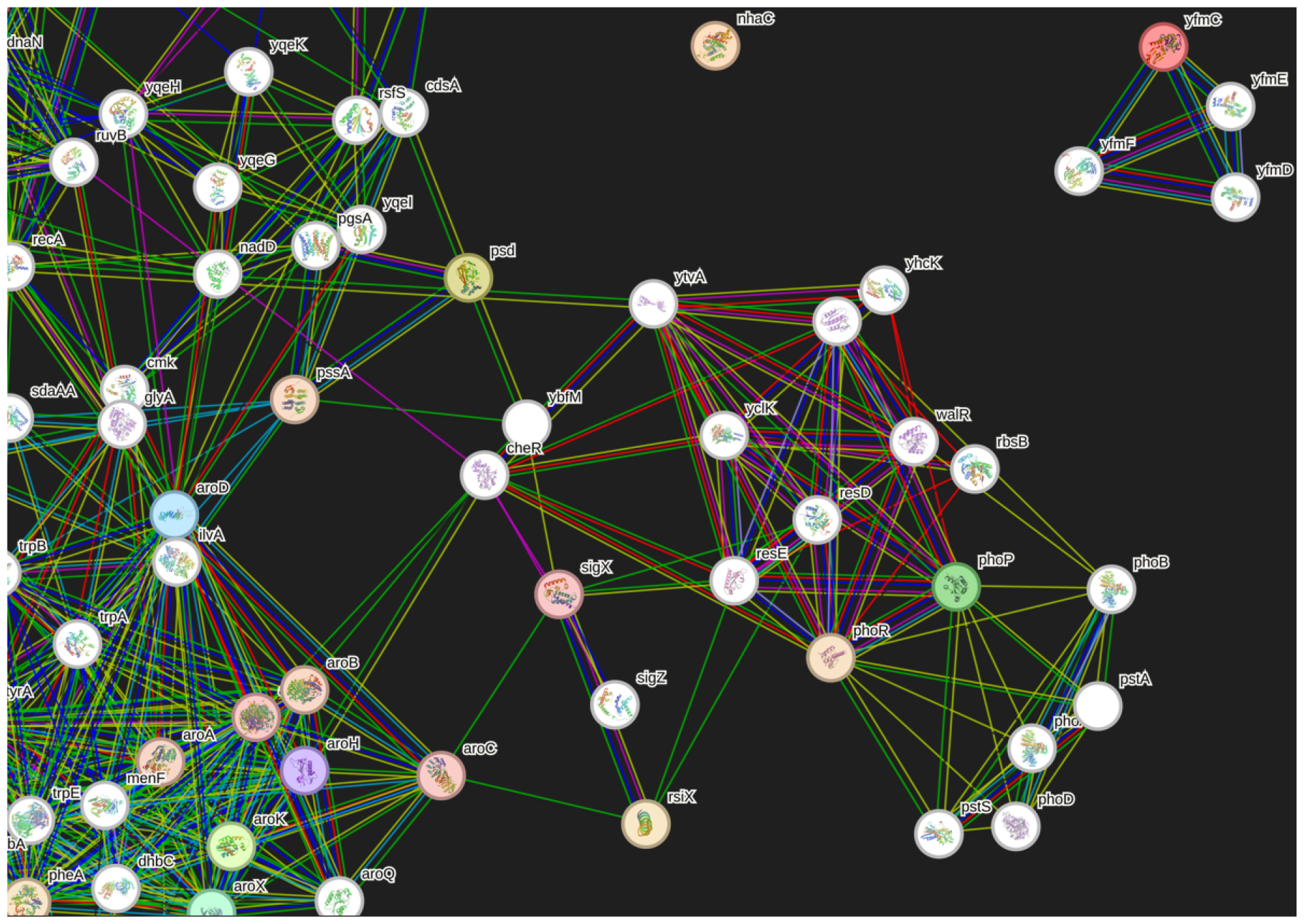
Portion of the genetic network connecting the arok family (Shikimate Kinase) to the yfmC family (ABC transport).Light blue and purple show known interactions from databases and experiments, respectively. Green, dark blue, and red lines signify predicted interactions (gene neighborhood, gene fusions, gene co-occurrence). Yellow lines represent text mining connections; black represents co-expression, and teal represents protein homology. There were two direct pathways found connecting the two protein families of interest. A) arok *→* SigX *→* resE *→* yclK *→* walk *→* yfmF *→* ymfC. B) arok *→* cheR *→* yclk *→* walk *→* yfmF *→* yfmC.

A confusion matrix was created for the general antibiotic predictive QSAR model to quantify the predictions’ accuracy.

Subsequently, another QSAR model was made to predict the testing set with known gram-positive antibiotics to relate to the mutation data that will be explored later. The training set consisted of 36 known gram-positive and 60 known non-gram-positive antibiotics. The same model development was used here. Lastly, it should be noted that, unlike the first QSAR trial, none of these compounds were eliminated from further computational testing, as there was a very significant probability that novel gram-positive inhibitors could be structurally dissimilar to current antibiotics, which would be reflected in the QSAR predictions.

Finally, following the same rationale as above and for thoroughness, the compounds were predicted to be gram-negative antibiotics. Unlike the two previous instances, the probability (p) was *≤* 0.5, where previously it had been 0.5 *≤ p ≤* 1. This means that the QSAR model did not predict any compound from the plant to be a gram-negative antibiotic.

### 3.3 Introduction of Shikimate Kinase

Following the comparative mutation analysis, numerous mutations were discovered throughout the three strains of mutant b.subtilis (mutant for roots, stems, and leaves). When these were compared with the parent strain, the common mutations throughout the three strains were noted as they would have a direct correlation towards the compound(s) that were contributing towards the antimicrobial effect of the goldenrod plant.

### 3.4 QSAR Comparison with Shikimate Kinase Inhibitors

Once shikimate kinase had been identified as the target/inhibitory mechanism of the antibiotic compound from the goldenrod plant, another QSAR was run against known (a small set) shikimate kinase inhibitors. These were all found in other literature. This QSAR also utilized a normal approach with a 70-30 split (random division).

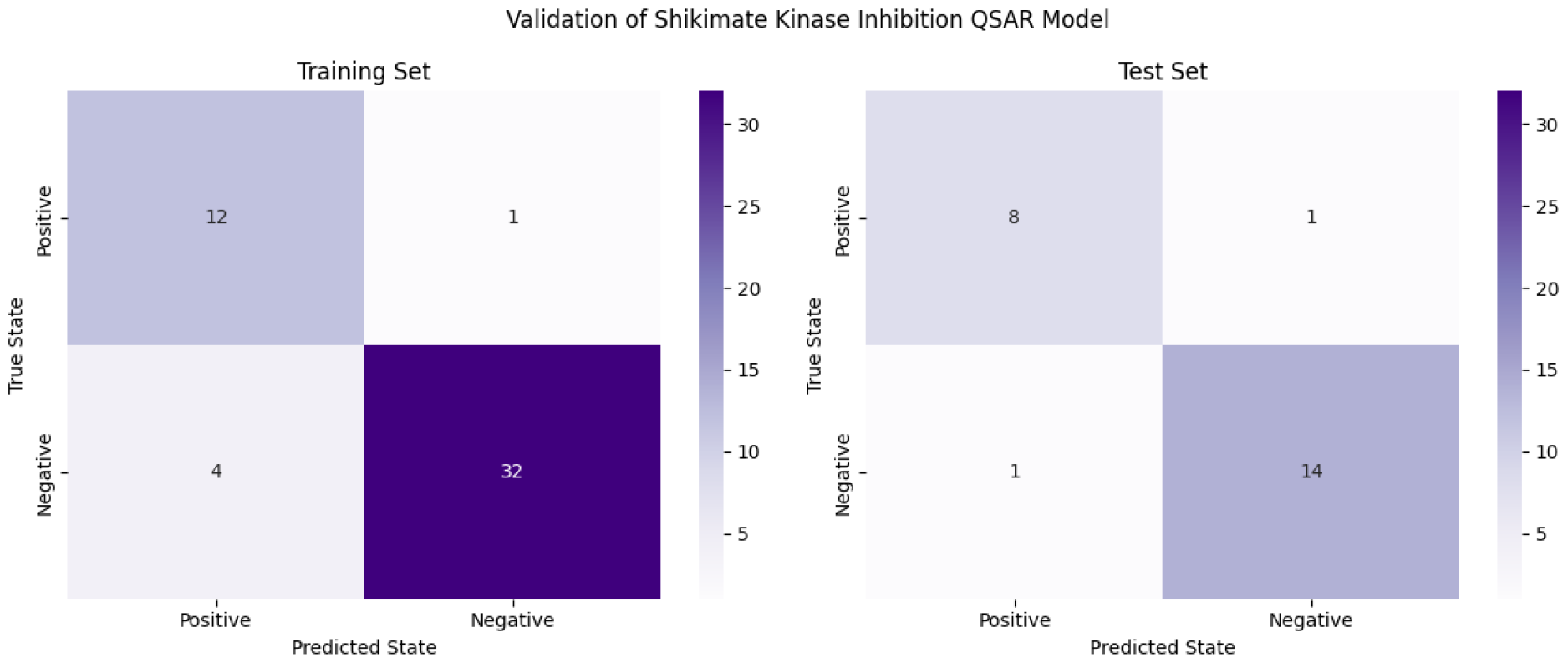
caption Confusion matrix for Shikimate Kinase Inhibition QSAR model

### 3.5 Molecular Docking

Out of the final five compounds that the QSAR model predicted, molecular docking simulations were run to determine binding affinity and, later, the inhibition constant. The equation for conversion following [1] is the following:

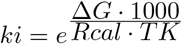

Where Δ*G* represents the binding affinity, Rcal is 1.98719, and TK is the temperature in kelvin (298.15).

Afterward, several post-docking programs were run to determine the interactions between the ligand and the pocket in the protein.

### 3.6 Post Molecular Docking Analysis

Subsequently, the most likely compound (C4) was chosen and was further analyzed to determine the specific interactions between Shikimate Kinase (Tables 7-9), and lastly, the pocket site for the ligand was analyzed to grade druggability to further validate the computational claims

**Table 1:** 12 Molecular Descriptors Chosen from Rdk Mordred 1826 [23] as well as a short description for each.

| Descriptor | Explanation |
| --- | --- |
| MW | Molecular weight of the compound. |
| nHeavyAtom | Number of heavy (non-hydrogen) atoms. |
| SLogP | LogP value computed from the simplified molecular-input line-entry system (SMILES). |
| SMR | Molar refractivity calculated from the SMILES. |
| nHBAcc | Number of hydrogen bond acceptors. |
| nHBDon | Number of hydrogen bond donors. |
| TopoPSA | Topological polar surface area. |
| nRot | Number of rotatable bonds. |
| naRing | Number of aromatic rings. |
| Lipinski | Lipinski’s rule of five, a rule of thumb to evaluate drug likeness. |
| FilterItLogS | Predicted aqueous solubility. |

**Table 2:** List of the top 22 predicted compounds from the testing set. All compounds predicted by the model having an antibiotic property probability from [0.5, 1] were chosen as possible antibiotics. Compounds listed starting with NP indicate that they are not available in any publicly available chemical data bank.

|  |  |  |  |  |  |
| --- | --- | --- | --- | --- | --- |
| C<sub>25</sub>H<sub>26</sub>O<sub>13</sub> | C<sub>17</sub>H<sub>20</sub>O<sub>9</sub> | C<sub>27</sub>H<sub>30</sub>O<sub>15</sub> | Np-02 | C<sub>6</sub>H<sub>6</sub>O<sub>3</sub> | C<sub>16</sub>H<sub>18</sub>O<sub>9</sub> |
| C<sub>8</sub>H<sub>9</sub>NO<sub>2</sub> | C<sub>18</sub>H<sub>16</sub>O<sub>8</sub> | C<sub>18</sub>H<sub>35</sub>NO | C<sub>15</sub>H<sub>12</sub>O<sub>6</sub> | NP-022035 | Np-000308 |
| C<sub>9</sub>H<sub>10</sub>O<sub>5</sub> | Np-03 | C<sub>22</sub>H<sub>30</sub>O<sub>14</sub> | C<sub>16</sub>H<sub>14</sub>O<sub>6</sub> | Np-09 | Np-000587 |
| NP-003692 | C<sub>15</sub>H<sub>10</sub>O<sub>5</sub> | C<sub>30</sub>H<sub>50</sub>O<sub>2</sub> | C<sub>15</sub>H<sub>20</sub>O<sub>4</sub> |  |  |

**Table 3:** List of 9 compounds predicted by the QSAR model to be gram-positive antibiotics, the predictive range was once again from [0.5, 1].

|  |  |  |  |  |  |
| --- | --- | --- | --- | --- | --- |
| NP-022035 | Np-09 | C<sub>21</sub>H<sub>20</sub>O<sub>11</sub> | C<sub>27</sub>H<sub>30</sub>O<sub>15</sub> | C<sub>18</sub>H<sub>31</sub>NO<sub>4</sub> | C<sub>6</sub>H<sub>6</sub>O<sub>3</sub> |
| C<sub>8</sub>H<sub>9</sub>NO<sub>2</sub> | C<sub>25</sub>H<sub>26</sub>O<sub>13</sub> | C<sub>22</sub>H<sub>30</sub>O<sub>14</sub> |  |  |  |

**Table 4:** Compounds with the highest probability of being gram-negative antibiotics, ranging from probabilities from 0.1 to 0.37.

|  |  |  |  |  |  |  |  |
| --- | --- | --- | --- | --- | --- | --- | --- |
| NP-022035 | C<sub>25</sub>H<sub>26</sub>O<sub>13</sub> | C<sub>8</sub>H<sub>9</sub>NO<sub>2</sub> | C<sub>22</sub>H<sub>30</sub>O<sub>14</sub> | C<sub>27</sub>H<sub>30</sub>O<sub>15</sub> | Np-09 | C<sub>6</sub>H<sub>6</sub>O<sub>3</sub> | C<sub>9</sub>H<sub>10</sub>O<sub>5</sub> |

**Table 5:** List of Proteins and Enzymes.

|  |  |
| --- | --- |
| Helicase<br>Integrase<br>2,4 Dienoyl CoA Reductase<br>Multidrug Resistance Efflux Transporter Family Protein<br>MFS Transporter<br>Shikimate Kinase<br>Phosphatidylserine Decarboxylase<br>Transcriptional Regulator PhoB | Na<sup>+</sup>/H<sup>+</sup><br>Prepilin Peptidase<br>tRNA (adenine-N(1)) - methyltransferase<br>RNA polymerase sigma factor Sig X<br>DEAD/DEAH Box Helicase<br>Sigma 54 dependent FIS family transcriptional regulator<br>HTH-type transcriptional repressor NsrR<br>ABC transporter ATP-binding protein |

**Table 6:** List of 5 compounds predicted by the QSAR model to be shikimate kinase inhibitors, the predictive range was once again from [0.5, 1].

|  |  |  |  |  |
| --- | --- | --- | --- | --- |
| C<sub>18</sub>H<sub>16</sub>O<sub>8</sub> | C<sub>15</sub>H<sub>10</sub>O<sub>5</sub> | C<sub>15</sub>H<sub>12</sub>O<sub>6</sub> | C<sub>11</sub>H<sub>9</sub>NO<sub>2</sub> | Np-000308 |

**Table 7:** Hydrophobic interactions between the ligand and receptor. The residue is the specific amino acid residue within the receptor protein. AA stands for an amino acid. The distance is measured in °A.

| Index | Residue | AA | Distance | Ligand Atom | Protein Atom |
| --- | --- | --- | --- | --- | --- |
| 1 | 118A | PRO | 3.79 | 9 | 1063 |
| 2 | 118A | PRO | 3.4 | 17 | 1062 |
| 3 | 119A | LEU | 3.56 | 13 | 1072 |

**Table 8:** Table showing the hydrogen bonding between the ligand and the receptor. The same notations in Table 7 are used.

| Index | Residue | AA | Distance H-A | Distance D-A | Donor Angle | Protein Donor | Side Chain | Donor Atom | Acceptor Atom |
| --- | --- | --- | --- | --- | --- | --- | --- | --- | --- |
| 1 | 54A | GLU | 2.57 | 3.18 | 121.21 | n | y | 18 [O3] | 488 [O.co2] |
| 2 | 58A | ARG | 3.15 | 3.87 | 128.78 | y | y | 533 [Ng+] | 2 [O2] |
| 3 | 58A | ARG | 2.03 | 3.03 | 167.32 | y | y | 534 [Ng+] | 2 [O2] |
| 4 | 81A | GLY | 2.38 | 3.09 | 125.49 | y | n | 736 [Nam] | 20 [O3] |
| 5 | 136A | ARG | 3.23 | 3.96 | 129.23 | y | y | 1243 [N3+] | 20 [O3] |
| 6 | 136A | ARG | 3.15 | 3.92 | 132.69 | y | y | 1245 [N3] | 18 [O3] |

**Table 9:** Pi Stacking Interactions in Molecular Docking. Details include interaction index, involved residues, pi stacking parameters (distance, angle, offset), type, and ligand atoms

| Index | Residue | AA | Distance | Angle | Offset | Stacking Type | Ligand Atoms |
| --- | --- | --- | --- | --- | --- | --- | --- |
| 1 | 49A | PHE | 3.71 | 9.45 | 0.37 | P | 12, 13, 14, 15, 16, 17 |

**Table 10:** Protein Ligand Profiler Data. Name represents pockets, Drug Score is a druggability score computed from various metrics with values ranging from 0 to 1. Pockets with scores *≥* 0.7 are good drug targets, while scores in the [0.86, 1] range represent phenomenal drug targets. A simple score is a score computed from 3 metrics: shape, size, and hydrophobicity.

| Name | Volume Å<sup>3</sup> | Surface Å<sup>2</sup> | Drug Score | Simple Score |
| --- | --- | --- | --- | --- |
| P_0 | 785.73 | 960.3 | 0.84 | 0.48 |
| P_1 | 154.05 | 294.65 | 0.34 | 0 |
| P_2 | 141.82 | 366.98 | 0.26 | 0 |
| P_3 | 113.15 | 294.53 | 0.22 | 0 |

## 4 Discussion

### 4.1 Preliminary Investigation

The discovery that the goldenrod plant exhibits inhibitory properties against B. subtilis and E. coli is corroborated by several studies. For instance, research published in the Journal of Agricultural and Food Chemistry (2021) demonstrates the antimicrobial potency of goldenrod root compounds against pathogens, including Bacillus subtilis. Similarly, a review of Solidago virgaurea L. underscores its traditional medicinal uses and highlights its antimicrobial potential. Further, the study on antimicrobial diterpenes from Solidago rugosa Mill. Strengthens this evidence by identifying specific compounds with antibacterial activity. These findings align with the observed effectiveness of Goldenrod in this study, suggesting that it may produce compounds with broad-spectrum antibacterial capabilities, warranting further investigation.

### 4.2 QSAR Experimentation

Identifying potential antibiotic compounds through QSAR analysis, based on their structural similarity to known antibiotics, is critical. However, this approach assumes that all effective antibiotics must share a structural resemblance with existing ones, potentially overlooking novel compounds with unique mechanisms. This limitation highlights the need for a more inclusive approach in future drug discovery efforts to ensure that structurally distinct yet effective compounds are not missed.

### 4.3 Shikimate Kinase Identification

The literature review highlights Shikimate Kinase as a crucial target in bacterial inhibition, aligning with findings related to plant-derived compounds, explicitly limiting the scope of plant-based compounds to affect cell wall synthesis and metabolic pathway inhibition [26, 27]. This specificity is notable, as it was observed that the inhibition of Shikimate Kinase, consistent with the action of other plantbased compounds, is pivotal in causing bacterial death. Furthermore, using the STRING database, the investigation into the potential interplay between the Shikimate Pathway and MFS/ABC transporters revealed some links [20]. However, these connections were not strong enough to challenge the hypothesis that Shikimate Kinase inhibition is the primary driver of bacterial mortality. Figure 13 below shows the complexity of the pathway connecting the upregulation of certain compounds/proteins in the Shikimate Acid pathway to their connection with MFS or ABC transport. Furthermore, many of these links are hypothetical and do not necessarily mean a specific connection, reducing the credibility of the alternative theory and ultimately supporting the simpler that shikimate kinase could mutate but still maintain functionality. Other studies have backed this [25], showing that non-functional shikimate kinase results in bacterial death for 99% of mutant strains.

### 4.4 Molecular Docking

The binding affinities observed for compounds C2 and C4 in this study, at -8.3 and -8.9 kcal/mol, respectively, are within the range of efficacious inhibitors reported in other studies using AutoDock Vina. These findings suggest that the identified compounds are potential inhibitors of Shikimate Kinase, offering promising avenues for drug development.

### 4.5 Post Molecular Docking Analysis

The molecular interactions observed, particularly the hydrogen bonds predominantly with arginine, are consistent with other studies on Shikimate Kinase inhibitors. The high druggability score of 0.84 for the receptor-binding pockets used by the ligands further substantiates their potential as effective drug candidates. The mechanism for inhibition is most likely competitive inhibition, as the pocket (active site) for both C4 and Shikimate Acid is the same, albeit with different residue binding. This suggests that although C4 binds to different residues because of its presence in the microenvironment, it acts as a competitive inhibitor.

## 5 Conclusion

Based on the data obtained in the study, it was shown that certain compounds in the goldenrod plant contribute chiefly to its antibiotic properties against gram-negative and gram-positive bacteria. Furthermore, the similarity of these compounds to other inhibitors of the target mechanism, Shikimate Kinase, was shown computationally. The compounds interact with many arginine residue chains and Leucine and phenylalanine residue chains. This research was tested through various computational techniques ranging from QSAR experimentation to Molecular Docking and protein-ligand binding analyses. The inhibition of Shikimate Kinase shows the potential for developing new spectrum antibiotics that are as potent as preexisting ones yet have lesser toxicity to mammalian cells. Additionally, creating derivations of this compound can allow for retained efficacy for significant periods. Future steps are to test these findings *in vitro* to gain a higher degree of confidence from the data collected *in sillico*.

## 6 Acknowledgements

I extend my deepest gratitude to Mr. Mathieu, who was pivotal in mentoring and steering this research journey. Profound thanks are also due to Professor Chai, whose expertise in mutation analysis and genomics significantly enriched the experimental scope of this work. Additionally, heartfelt appreciation is extended to Professor Sjoelund, whose generous contribution to LS-MS spectrometry services was instrumental in advancing the analytical aspects of this study.

